# Bridging Prediction and Reality: Comprehensive Analysis of Experimental and AlphaFold 2 Full-Length Nuclear Receptor Structures

**DOI:** 10.1101/2025.03.01.640955

**Authors:** Akerke Mazhibiyeva, Tri T. Pham, Karina Pats, Martin Lukac, Ferdinand Molnár

**Affiliations:** Laboratories of Computational Structural Biology, Nazarbayev University, Kabanbay Batyr 53, Astana, 010000, Kazakhstan; Laboratories of Mechanobiology, Department of Biology, Nazarbayev University, Kabanbay Batyr 53, Astana, 010000, Kazakhstan; Institute of Applied Computer Sciences, ITMO University, 49, Kronverksky Ave., 197101, Saint Petersburg, Russian Federation; Department of Computer and Networks Engineering, Hiroshima City University, 3-4-1 Oozukahigashi, Asaminami-ku, Hiroshima-shi, 7313166 Japan

## Abstract

AlphaFold 2 has revolutionized protein structure prediction, yet systematic evaluations of its performance against experimental structures for specific protein families remain limited. Here we present the first comprehensive analysis comparing AlphaFold 2-predicted and experimental nuclear receptor structures, examining root-mean-square deviations, secondary structure elements, domain organization, and ligand-binding pocket geometry. While AlphaFold2 achieves high accuracy in predicting stable conformations with proper stereochemistry, it shows limitations in capturing the full spectrum of biologically relevant states, particularly in flexible regions and ligand-binding pockets. Statistical analysis reveals significant domain-specific variations, with ligand-binding domains showing higher structural variability (CV = 29.3%) compared to DNA-binding domains (CV = 17.7%). Notably, Alphafold 2 systematically underestimates ligand-binding pocket volumes and captures only single conformational states in homodimeric receptors where experimental structures show functionally important asymmetry. These findings provide critical insights for structure-based drug design targeting nuclear receptors and establish a framework for evaluating Alphafold 2 predictions across other protein families.

## INTRODUCTION

As of January 2025, the Protein Data Bank, hosted at rcsb.org (RCSB PDB), contained 230,083 experimental biological macromolecular structures ^1,2^, whereas the UniProtKB/TrEMBL protein database ([release 2024_06](https://www.ebi.ac.uk/uniprot/TrEMBLstats)) contained 254,254,987 sequence entries ^3^. The structural gap in structural genomics refers to a state where a generated data from the next and third generation sequencing and the subsequent *in silico* prediction of protein-coding sequences grows over the time faster than the number of experimentally determined protein structures. By researchers in the structural biology community, the proposed solution to bridge this gap is the AlphaFold 2 (AF2) system, which combines computational algorithms and artificial intelligence to efficiently predict and build structural models ^4–7^. All the AF2 predicted structures can be obtained from AlphaFold Protein Structure Database (https://alphafold.ebi.ac.uk/) and recently also from RCSB PDB ^1^. For training, the AF2 algorithm has been using protein structures that are mostly from the PDB release prior to April 30, 2018 with the addition of some structures obtained from the PDB release before February 15, 2021^4,6^. However, it has been reported that PDB templates are not required for an accurate AF2 predictions, as it can produce *de novo* models using multiple sequence alignments (MSA) alone. Moreover, AF2 may also disregard the PDB templates with low quality ^5^. Although AF2 is incapable of predicting accurately the positions of cofactors, metals, ions, and nucleic acids, which are found in many experimental structures, it is trained to predict protein structures as close to their native conformation as possible ^4^. Therefore, the positions of the backbone and side chains are usually consistent with the expected structures that contain cofactors or ions. Nevertheless, the absence of additional structural information may result in an inaccurate or improper prediction of the protein structure ^6^. The AF2 models accuracy might vary both within and between structures depending on the particular protein ^4^. This accuracy can be assessed by the predicted local distance difference test (pLDDT) score which is provided for each residue ^5^. Nonetheless, a specific problem manifests in case of disordered protein regions that produce a low-confidence pLDDT score below 50. The <50 score suggests that the region is unstructured in a physiological environment or requires an additional interaction partner(s) such found in native protein complexes. Indeed, many proteins show variable degrees of intrinsic disorder and thus require additional partners and it has been estimated that at least 30% of the human proteome are intrinsically disordered proteins ^8^. In addition, there is a significant amount of proteins that contain multiple domains. These two features bring high complexity to protein structures located in complexes. The regions with a score between 50 and 70 are likewise poorly modeled and have low confidence whereas range between 70 and 90 indicate better models producing a good backbone prediction. Finally, regions with a score higher than 90 are expected to have the highest accuracy ^5^.

Based on our expertise and the importance of the nuclear receptor (NR) superfamily as drug targets, we have chosen to evaluate their AF2 predicted and experimental structures. In particular, full-length (FL) multi-domain NRs are of importance since most of them are not present in the PDB. AF2 might be very useful for NR research, specifically to understand the structure-function relationship for their potential as therapeutic targets. For this purpose, it is important to assess the reliability of NR models predicted by AF2. NRs are ligand-activated transcription factors that are responsible for sensing metabolic and systemic hormonal signals ^9^. Once activated, they regulate the transcription of genes that control a wide range of cellular processes including cell proliferation, growth, differentiation, and metabolism. Human NRs can be divided into three subgroups based on the nature and affinity of their ligands: i) endocrine steroid hormone receptors, ii) nutritional sensors (adopted orphans) and iii) orphan receptors ^10^. Endocrine receptors were identified when researchers were looking for the receptors of well-known steroids such as testosterone, estradiol, progesterone, cortisol and aldosterone. Orphan receptors are NRs for which until now no ligand has been identified or no ligand exists for them. However, earlier, for some of the orphans natural or xenobiotic compounds were identified as their ligands and thus, these receptors became adopted orphan receptors. NRs are involved in nearly all human cellular processes, thus they are one of the most established and studied drug targets, being responsible for the therapeutic effect of 16% of small-molecule drugs ^11,12^.

Given the importance of NRs as drug targets and the scarcity of FL multi-domain NR structures in the PDB, we conducted the first comprehensive assessment comparing AF2-predicted conformations to all accessible experimental three-dimensional (3D) structures, providing a critical benchmark to guide future structural and drug discovery efforts in this important protein family.

## RESULTS

### The availability of structural data for human NRs in PDB

To date (January 2025), the 12 human steroid hormone NRs have altogether 817 solved structures, the estrogen receptor (ER) *α* being the most studied with 454 structures yet from all of them only retinoic acid receptor (RAR) *β* and glucocorticoid receptor (GR) have a FL multi-domain structural data available. From the 27 human nutritional sensors there are 1026 accessible structures in the PDB, the proliferator-activated receptor (PPAR) *γ* having the highest number e.g. 316 structures available in single and as FL multi-domain 3D structures. Besides PPAR*γ*, there are FL multi-domain 3D structures for hepatocyte nuclear factor (HNF)4*α*, liver X receptor (LXR) *β*, NR related 1 protein (NURR1) and retinoid X receptor (RXR) *α*. Two NRs from this family the RAR-related orphan receptor (ROR) *β* and testicular NR (TR) 2 lack any kind of structural data. The 9 human orphan NRs have the lowest presence in the PDB mostly in the form of peptides e.g. 20 structures. The germ cell nuclear factor (GCNF) does not have any available 3D structures in PDB. Out of 48 human NRs altogether seven human FL multi-domain NR 3D structures are available in the PDB (Table 1): i) two GR ^13^ ii) one (HNF4*α*) homodimer ^14^, iii) one RXR*α*-LXR*β* heterodimer ^15^, iv) one NURR1^16^ v) three PPAR*γ*-RXR*α* heterodimers ^17^, vi) one RAR*β*-RXR*α* heterodimer, and vii) the FL RXR*α* present in all three heterodimers. ^18^.

**Table 1.** Summary of metrics comparing experimental and AlphaFold 2 NR structures.

| Metric | Experimental Structures | AF2 Predictions | Significant Differences | Biological/Functional Implications |
| --- | --- | --- | --- | --- |
| RMSD (by domain) | Variable structural similarity between domains | Generally low RMSD values (<0.55 Å) | LBDs showed higher structural variability (CV = 29.3%) than DBDs (CV = 17.7%); DBD RMSD exceeded LBD in 41.7% of cases | Flexible regions essential for ligand binding have higher prediction variability |
| SSE Content | Higher loop content, variable α-helical distribution | Tendency toward higher α-helical content (+4.2% DBDs, +3.8% LBDs) with compensatory decreases in loop regions (-5.0%) | Strong negative correlations between α-helical and loop content ($r = -0.527$ for DBD, $r = -0.906$ for LBD) | AF2 may overestimate secondary structure stability in dynamically flexible regions |
| Domain Distances ($\epsilon_{\text{DBD,LBD}}$) | Marked asymmetry in homodimeric structures; partner-specific distances in heterodimers | Captures only a single conformational state | Homodimers show bimodal distance distribution not captured by AF2 | AF2 misses functionally important asymmetry in domain organization |
| Domain Angles ($\theta_{\text{DBD,LBD}}$) | Receptor-specific and partner-dependent variations | Generally predicts one angle conformation | RXRα showed most partner-dependent variations not fully captured by AF2 | Angle variations may reflect allosteric regulation mechanisms |
| Dihedral Angles ($\text{DH}_{\text{DBD,LBD}}$) | Distinct conformational clusters, particularly in NURR1 and GR | Captures predominant conformational states | GR showed bimodal distribution (IQR = 81.81°) not fully captured by AF2 | Dihedral angle diversity may be essential for regulatory element interactions |
| LBP Geometry | Larger, more variable pocket volumes | Systematically underpredicts pocket volumes (by 8.4% on average) | LXRβ showed most significant volume difference (>150% larger in experimental structures) | May impact structure-based drug design, potentially missing cryptic pockets |
| Structure Validation Parameters | Variable RSCC scores, especially in flexible regions | Consistent pLDDT patterns with low scores in intrinsically disordered regions | Low pLDDT scores correlate with missing residues or low RSCC values | Highlights regions involved in conformational changes and allosteric regulation |
| Backbone Torsional Analysis | More Ramachandran outliers, often in functionally relevant regions | Higher stereochemical quality with fewer outliers | Experimental PPARγ structure had 18 Ramachandran outliers vs. only 1 in AF2 | "Imperfections" in crystal structures may represent legitimate functional states |

### Root-mean-square deviation of the structures

To assess the fine differences between the protein structures, three root-mean-square deviation (RMSDs) metrics have been calculated: i) all-atoms (Fig.1A column ***a***), ii) backbone (N, C*α*, C and O atoms) (Fig.1A column ***b***) and iii) only C*α* carbons (Fig.1A column ***c***). This comprehensive approach allowed us to overcome the positional domain bias that was addressed later in the “Geometric analysis of domain architecture section” and is due to differential positioning of predicted and experimental protein domains in the 3D space. Statistical analysis revealed distinct patterns in structural deviation, with mean RMSD values for DNA-binding domains (DBDs) of 0.442 ± 0.078 Å (all-atom), 0.379 ± 0.077 Å (backbone), and 0.359 ± 0.080 Å (C*α*), while ligand-binding domains (LBDs) showed slightly higher deviations of 0.512 ± 0.150 Å (all-atom), 0.452 ± 0.135 Å (backbone), and 0.442 ± 0.137 Å (C*α*). The AF2 predicted models for the majority of NRs demonstrated high structural similarity with experimental structures, showing all-atom RMSD less than 0.55 Å for both DBD and LBD domains (Fig.1A). Notable exceptions were observed in GR LBDs (PDBID: 7PRW both chain A and B), NURR1 LBDs (PDBID: 7WNH all chains A, B, C, D), and LXR*β* DBD (PDBID: 4NQA chain I). Quantitative assessment revealed significant domain-specific differences (mean difference = -0.070 Å, t = -1.988, p < 0.05, Cohen’s *d* = 0.574), with DBD RMSD values unexpectedly exceeding those of LBD in 41.7% of cases (10 out of 24 structures). The conventional hierarchical pattern of RMSD values (all-atom > backbone > C*α*) was not maintained in 12.5% of the analyzed structures, while the backbone and C*α* RMSDs remained consistently below 0.50 Å, except for the aforementioned NURR1 and LXR*β* structures. Importantly, individual NR analysis revealed receptor-specific variation patterns: NURR1 showed the most extreme domain-specific differences (DBD: CV = 0.7%; LBD: CV = 18.0%, range = 0.379 Å), GR demonstrated moderate variation (DBD: CV = 3.1%; LBD: CV = 9.5%), while HNF4A uniquely showed higher variability in DBD predictions (CV = 14.7%) compared to LBD (CV = 7.9%). Overall, the LBD showed particularly higher structural variability (CV = 29.3%) compared to DBD (CV = 17.7%), suggesting distinct challenges in structural prediction across different NR families and their domains.

**Figure 1:**
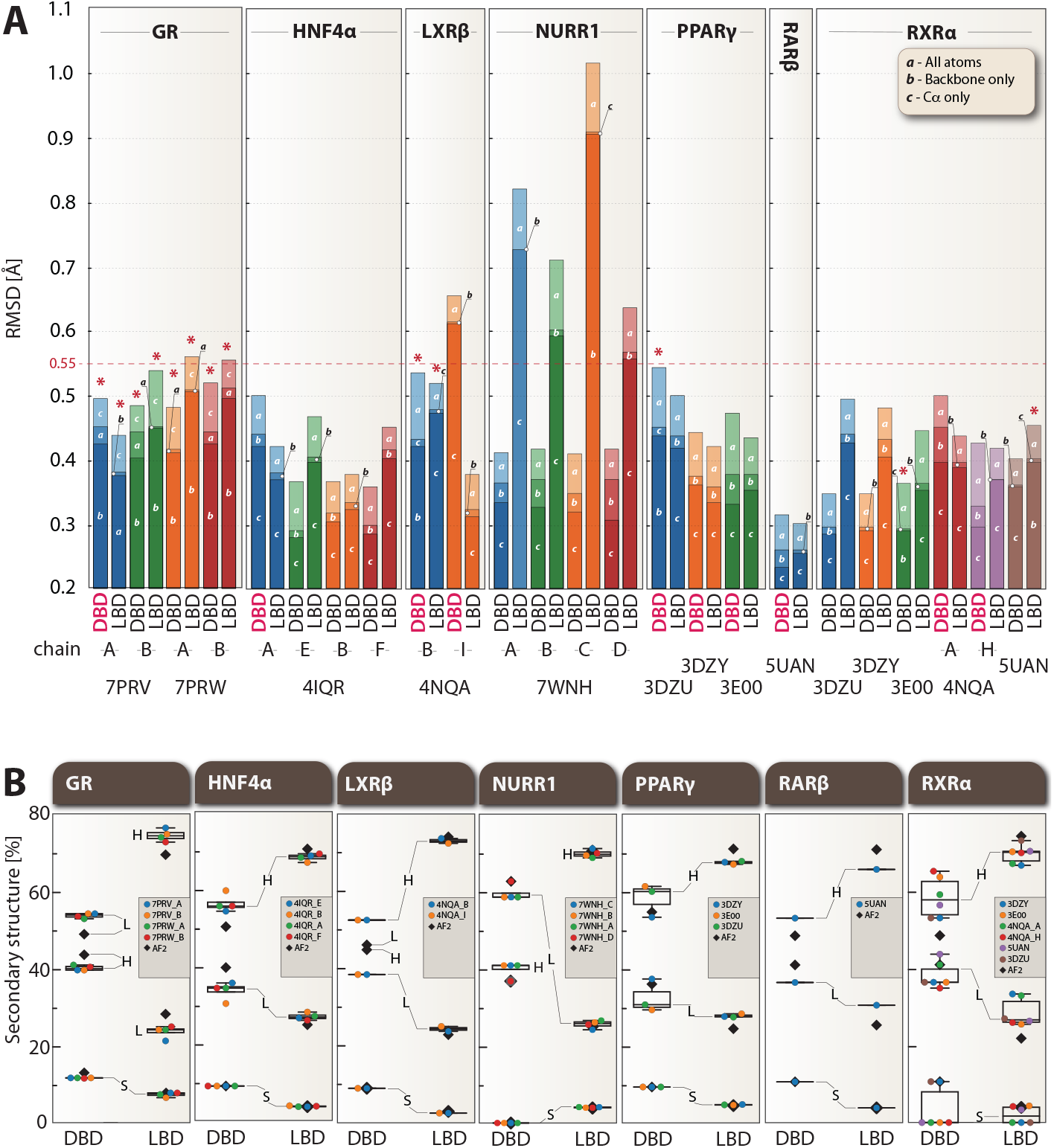
Analysis of RMSD and SSE content for experimental and AF2-predicted full-length NRs. A) RMSD values are shown for all atoms (*a*), backbone atoms only (*b*), and C*α* atoms (*c*), measured in Å. The columns are overlaid, and different color hues represent the conditions used to calculate RMSD. Exceptions to the expected hierarchical pattern (all-atom > backbone only > C*α* only) are marked with asterisks (*). Additionally, the 10 cases where the DNA-binding domain (DBD) shows higher RMSD than the ligand-binding domain (LBD) are highlighted with “DBD” in red text. B) Percentage distribution of secondary structure elements (SSEs) for *α*-helices (H), *β*-sheets (S), and loops (L) across all analyzed structures. In both panels, color coding distinguishes different chains: blue (first chain), orange (second chain), green (third chain), red (fourth chain), purple (fifth chain), brown (sixth chain), and gray (AF2 predictions, only in panel B).

### Secondary structure element content analysis

Secondary structure element (SSE) analysis was performed to assess the similarity of experimental and predicted structures in terms of the *α*-helical, *β*-sheet and loop content (Fig. 1B). Statistical analysis revealed systematic differences in SSE distribution, with AF2 predictions showing a general tendency toward higher *α*-helical content (mean difference = +4.2% ± 2.1% for DBDs, +3.8% ± 1.9% for LBDs, p < 0.05) and compensatory decreases in loop regions (−5.0% ± 2.3%, p < 0.05). *β*-sheet content showed minimal variation across structures (mean difference = +0.8% ± 0.5%). This pattern was particularly evident in individual NR analysis: GR DBD showed 3-4% more *α*-helical and 1.5% more *β*-sheet content in the predicted structure, while its LBD exhibited 4-7% lower *α*-helical proportion compared to experimental structures. HNF4*α* predictions displayed 5-10% higher DBD *α*-helical content, with preserved *β*-sheet proportions and reduced loop content. LXR*β* demonstrated similar trends in its DBD (6-7% higher *α*-helical content), while maintaining consistent LBD secondary structure patterns. NURR1 showed an inverse trend, with experimental structures (7WNH chain A, B, C) having approximately 4% higher DBD *α*-helical content compared to the AF2 structure. PPAR*γ* and RAR*β* predictions showed 3-5% higher *α*-helical content in both domains, with corresponding reductions in loop regions. RXR*α* displayed the most notable differences, with AF2 predictions showing 4-8% higher *α*-helical content and the presence of *β*-sheet structure (approximately 10% in DBD, 4% in LBD) where several experimental structures showed none. Strong negative correlations were observed between *α*-helical and loop content (r = -0.527 for DBD, r = -0.906 for LBD), indicating compensatory relationships between these SSEs.

### Geometrical analysis of domain architecture

#### Distance *ϵ*_*DBD,LBD*_ between DBD and LBD

The distance between the two center of masses (COMs) DBD_*COM*_ and LBD_*COM*_ was measured in order to assess the similarity of the experimental and predicted structures in terms of their DBD and LBD position and are displayed in Fig. 2B. Experimental GR PDBs consist of GR homodimer-DNA-peptide in complex with furoate (7PRV) and velsecorat (7PRW). The AF2 GR (42.664 Å) is more similar to 7PRV_B and 7PRW_B, which have DBD and LBD at a distance of 46.865 Å and 44.134 Å, respectively. 7PRV_A and 7PRW_A have a less distanced domains, both around 32 Å. The PDB 4IQR of experimental HNF4*α* consists of an asymmetric unit with two independent HNF4*α* homodimer-DNA-peptide complexes. The LBDs of NRs are symmetrical when superimposed separately, but the overall NR has an asymmetrical organization in a complex with DNA. The AF2 model at 65.058 Å is more similar to experimental HNF4*α* (4IQR_B, _F) which have DBD and LBD at a distance of around 66 Å at their COMs. The other experimental HNF4*α* (4IQR_A, _E) have less distanced domains, around 38-39 Å. For LXR*β*, it is seen that the relative positions of NR DBDs and LBDs slightly differ within the asymmetric unit of RXR*α*-LXR*β* heterodimers. Compared with the experimental structures, the LXR*β* AF2 model at 43.928 Å is more similar to 4NQA_I, which has a distance of 43.537 Å between its DBD and LBD COMs, than 4NQA_B with 45.331 Å. The NURR1 PDB (7WNH) consists of four monomer-DNA complexes. The AF2 model of NURR1 has DBD and LBD at a distance of 44.956 Å, which is more than experimental structures (7WNH_A at 37.72 Å; _B at 36.93 Å), but less than 7WNH_C 63.191 Å and _D 65.044 Å. PPAR*γ* has a closely positioned DBD and LBD both in experimental (3DZU: 38.657 Å; 3DZY: 39.162 Å; 3E00: 39.368 Å) and predicted model 39.343 Å. RAR*β* has less distanced DBD and LBD, the distance between DBD_*COM*_ and LBD_*COM*_ being slightly more than 35 Å for both experimental 5UAN 35.374 Å and predicted 35.143 Å structures. The DBD and LBD of RXR*α* partnered with LXR*β* (4NQA_A, _H) and RAR*β* (5UAN) are spatially displaced from each other, the distance between their COMs being around 54-58 Å. The distance is smaller for RXR*α* partnered with PPAR*γ* (3DZY, 3DZU and 3E00), around 46-47 Å. This value is also more close to the predicted model with 48.139 Å. A striking pattern emerged in the analysis of homodimeric structures (GR and HNF4*α*): both receptors display marked asymmetry in their domain organization when bound to DNA. In GR structures, one monomer exhibits a compact conformation (domains ∼32 Å apart in 7PRV_A/7PRW_A) while its partner adopts an extended state (∼44-47 Å in 7PRV_B/7PRW_B). Similarly, HNF4*α* shows asymmetric organization with one set of monomers having closer domain proximity (∼38-39 Å in 4IQR_A/E) compared to their partners (∼66 Å in 4IQR_B/F). Notably, AF2 predictions capture only a single conformational state for each receptor (GR: 42.664 Å; HNF4*α*: 65.058 Å), corresponding more closely to the extended conformations observed in experimental structures.

**Figure 2:**
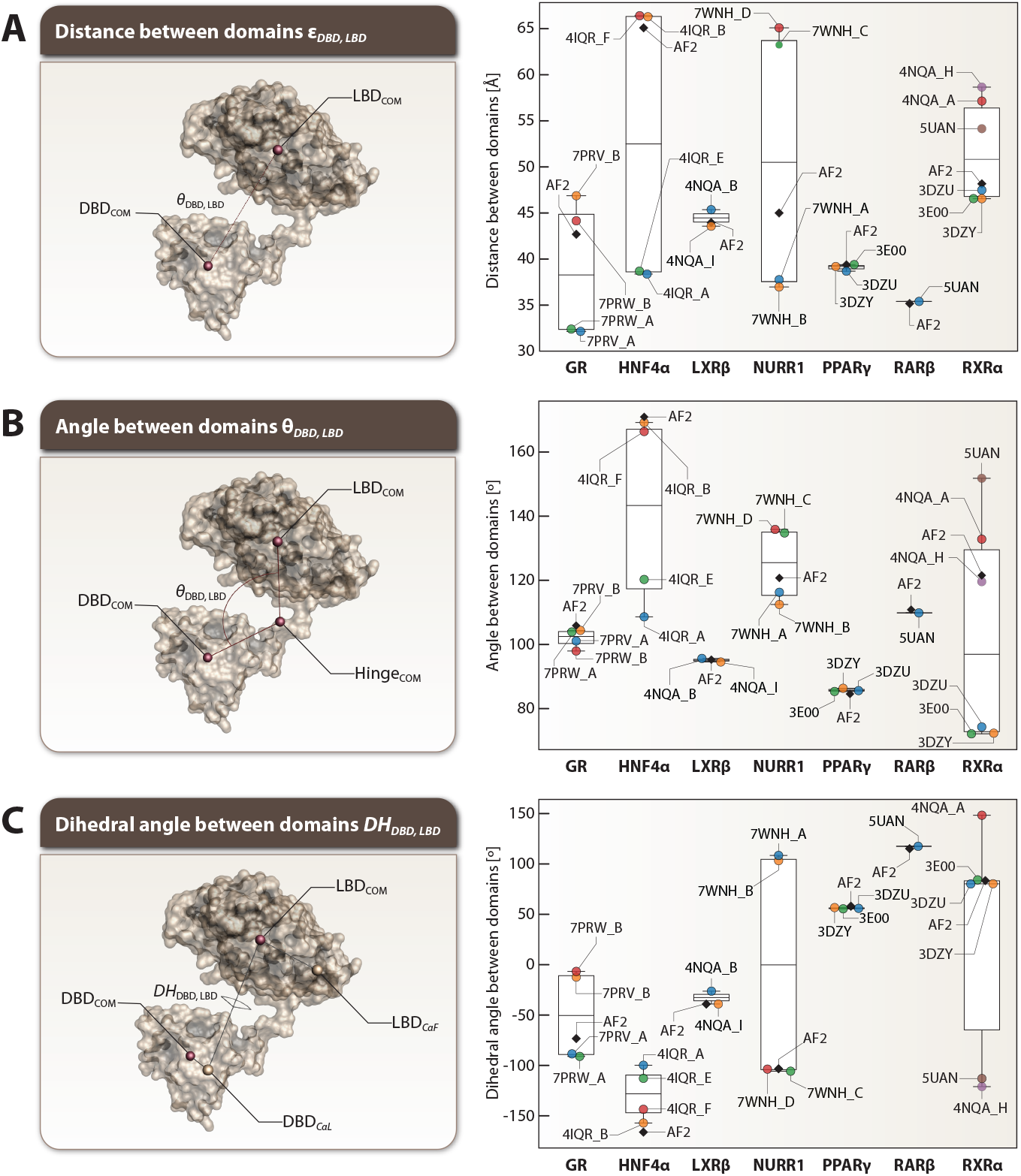
Domain organization of FL NRs analyzed through geometric measurements. A) Distances (*ϵ*_*DBD,LBD*_) between the COM of the DBD and LBD, reported in Å. Color coding distinguishes different chains: blue (first chain), orange (second chain), green (third chain), red (fourth chain), purple (fifth chain), brown (sixth chain), and gray (AF2 predictions). Notable patterns include marked asymmetry in homodimers: GR shows one monomer with compact conformation (∼ 32 Å in 7PRV_A/7PRW_A) and its partner in extended state (∼ 44-47 Å in 7PRV_B/7PRW_B); similarly, HNF4*α* displays asymmetric organization (38-39 Å in 4IQR_A/E vs. 66 Å in 4IQR_B/F). AF2 models predict only single conformational states (GR: 42.664 Å; HNF4*α*: 65.058 Å) corresponding to extended conformations. Partner-specific variations are observed in RXR*α* (46-47 Å with PPAR*γ* vs. 54-58 Å with LXR*β*/RAR*β*). B) Angles (*θ*_*DBD,LBD*_) formed by the COMs of the DBD, Hinge, and LBD, given in °. Significant variations include HNF4*α* showing two distinct angle conformations (108-120° in 4IQR_A/E vs. 166-169° in 4IQR_B/F); NURR1 exhibiting variable angles across structures (112-115° in 7WNH_A/B, 134-135° in 7WNH_C/D); and RXR*α* displaying partner-dependent angles (72-74° with PPAR*γ*, 119-132° with LXR*β*, 151° with RAR*β*). C) Dihedral Angles (*DH*_*DBD,LBD*_) defined by the COM of the DBD, the last C*α* atom of the DBD, the COM of the LBD, and the first C*α* atom of the LBD, reported in °. GR and NURR1 show bimodal distributions: GR varies between -91° to -88° (7PRV_A/7PRW_A) and -13° to -7° (7PRV_B/7PRW_B); NURR1 shows positive angles (103-108° in 7WNH_A/B) and negative angles (−106° to -103° in 7WNH_C/D). AF2 predictions generally capture predominant conformational states observed in experimental structures.

#### Angle *θ*_*DBD,LBD*_ between DBD and LBD

The angles between the DBD_*COM*_, Hinge_*COM*_ and LBD_*COM*_ were measured to specifically evaluate the relative positions of the DBD and LBD in experimental and predicted structures of NRs (Fig. 2B). GR *θ*_*DBD,LBD*_ is similar across experimental structures 7PRV_B 104°, 7PRW_A 103.528° and AF2 model 105.517°, but slightly differ for 7PRV_A 100.707° and 7PRW_B 97.590°. For HNF4*α*, the *θ*_*DBD,LBD*_ of two chains 4IQR_A 108° and 4IQR_E 120° is notably different from that of the AF2 model with 171°, while the two other chains 4IQR_B 169° and 4IQR_F 166° have angles similar to the predicted structure. LXR*β θ*_*DBD,LBD*_ is nearly identical across all analyzed structures at approximately 94-95°. As for NURR1, angles vary across the structures, with 7WNH_C 134.403° and 7WNH_D with 135.500° showing large values of *θ*_*DBD,LBD*_, while predicted structure has an angle of 120.367°. 7WNH_A 115.893° and 7WNH_B 112.096° show smaller *θ*_*DBD,LBD*_. PPAR*γ θ*_*DBD,LBD*_ is similar for the experimental 3DZU 85°, 3DZY 86° and 3E00 85° and predicted structure with 84°. For RAR*β*, the *θ*_*DBD,LBD*_ difference between experimental with 109° and predicted model with 110° is not significant. Experimental structures of RXR*α* partnered with PPAR*γ* (3DZU 74°; 3DZY 74° and 3E00 72°) have similar angles within each other; however, they considerably deviate from the predicted structure, which is at 121°. The predicted RXR*α θ*_*DBD,LBD*_ is most similar to that of RXR*α* partnered with LXR*β* 4NQA_A 132° and 4NQA_H 119°. RXR*α* from RAR*β*-RXR*α* heterodimer 5UAN has *θ*_*DBD,LBD*_ of 151°.

#### Dihedral angles *DH*_*DBD,LBD*_ between DBD and LBD

The dihedral angles between DBD_*COM*_, DBD_*CαL*_, LBD_*CαF*_ and LBD_*COM*_ were measured to evaluate the relative rotational relationship of the domains (Fig. 2C). The *DH*_*DBD,LBD*_ of experimental GR vary within each PDB. 7PRV_A with -88.633° and 7PRW_A with -91.144° have more negative values, while the other two chains 7PRV_B -12.502° and 7PRW_B -6.825° have dihedral angle closer to 0°. The *DH*_*DBD,LBD*_ of the predicted GR model is -73.475°. For all experimental HNF4*α* structures, the *DH*_*DBD,LBD*_ is negative, indicating that it is rotated counterclockwise. However, there is a big difference in values between two of the 4IQR chains (4IQR_A -100° and 4IQR_E -113°) and the AF2 model with -166°. The *DH*_*DBD,LBD*_ of the predicted structure is more similar to other two 4IQR chains (4IQR_B -157° and 4IQR_F -144°). The LXR*β* of 4NQA_I and the AF2 model have a *DH*_*DBD,LBD*_ of -39°, while 4NQA_B LXR*β* has -26°. Out of four NURR1 experimental structures, two chains 7WNH_A 108.236° and 7WNH_B 103.090° have positive dihedral angles, while the other two chains 7WNH_C -105.849° and 7WNH_D-103.796° along with the predicted NURR1 structure with -103.432°, have negative angles of similar values. Experimental 3DZU 56°; 3DZY 57° and 3E00 55° and predicted PPAR*γ* of 58° structures are similar in terms of their *DH*_*DBD,LBD*_, all being in a range of 55-58°. Similarly, there is no significant difference between the experimental 117° and predicted model with 115° of RAR*β DH*_*DBD,LBD*_. Experimental RXR*α* partnered with PPAR*γ* 3DZU 80°, 3DZY 80° and 3E00 84°, and AF2 model of 83° have a positive *DH*_*DBD,LBD*_ in the range of 80-84°. Although RXR*α* from 4NQA_B also has a positive angle, the value is substantially different, 148°. RXR*α* of 4NQA_I and 5UAN have a negative *DH*_*DBD,LBD*_ around -121°, and -113°, respectively.

### Analysis of ligand-binding pocket geometry

The analysis of ligand-binding pockets (LBPs) reveals distinct patterns between experimental structures and AF2 predictions (Fig. 3). Volumetric calculations show varying degrees of difference between experimental and predicted structures (Fig. 3A). GR structures 7PRV_A/B closely match AF2 predictions (∼3% difference), while 7PRW_A/B show 16-18% larger volumes. HNF4*α* experimental structures consistently show larger volumes than predicted 111.6-120%. LXR*β* displays the most significant volume differences, with experimental structures 4NQA_B and 4NQA_I exceeding AF2 predictions by 163.5% and 155.8%, respectively. NURR1, being a true orphan NR, exhibits an occluded LBP due to bulky aromatic residues in its *apo* form. PPAR*γ* shows mixed results, with 3DZU_D having 113.3% larger volume than AF2, while 3DZY_D and 3E00_D show smaller volumes (90.4% and 89.7%). RAR*β* experimental structure (5UAN_B) closely matches AF2 predictions (104.2%). RXR*α* structures show variable agreement, with 3E00_A, 4NQA_H, and 5UAN_A closely matching AF2 (≤3% difference), while 3DZU_A, 3DZY_A, and 4NQA_A show larger volumes (107.6-117.2%).

**Figure 3:**
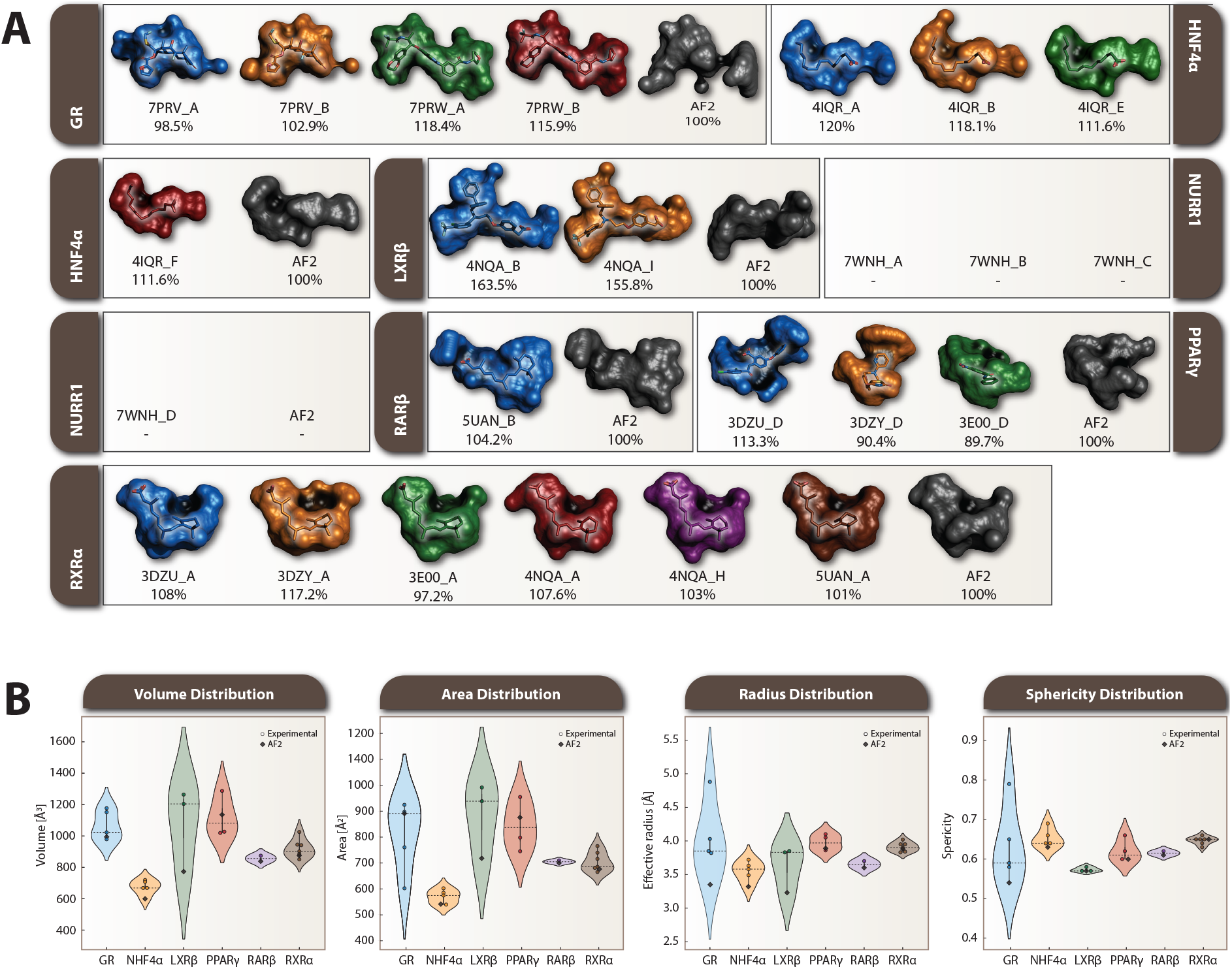
LBP metrics analysis across analyzed NRs. A) Surface representation of the calculated LBPs from FL NRs. The LBP of the AF2-predicted structure was set as 100% and served as a reference for normalizing the LBPs of other structures. Color coding distinguishes different chains: blue (first chain), orange (second chain), green (third chain), red (fourth chain), purple (fifth chain), brown (sixth chain), and gray (AF2 predictions). Notable variations include LXR*β* experimental structures exceeding AF2 predictions by >150%, while GR structures 7PRV_A/B closely match AF2 predictions (∼ 3% difference). NURR1 shows occluded LBPs in both experimental and predicted structures due to its orphan receptor status. B) Distribution of calculated LBP features: volume, area, effective radius, and sphericity across receptors. Statistical analysis reveals AF2 accurately predicts pocket sphericity (average deviation 3.2%, r=0.891) but systematically underpredicts effective pocket radius (average deviation 8.4%, r=0.643). A negative correlation (r=-0.65) between radius and sphericity indicates larger pockets tend to be less spherical. RXR*α* shows the most consistent predictions across parameters, while LXR*β* exhibits the largest radius discrepancy (−15.8%) and GR shows the highest sphericity variation (−7.9%).

Statistical analysis using Kruskal-Wallis tests confirms significant differences between receptors in both effective radius (H=32.145) and sphericity (H=27.893). AF2 demonstrates high accuracy in predicting pocket sphericity with average deviation 3.2%, slope=0.967, r=0.891 but systematically underpredicts effective pocket radius with average deviation 8.4%, slope=0.892, r=0.643. A negative correlation between effective radius and sphericity r=-0.65 suggests larger pockets tend to be less spherical. RXR*α* shows the most consistent predictions across both parameters effective with radius deviation -0.5%, sphericity deviation 0.0%, while LXR*β* exhibits the largest radius discrepancy of -15.8% and GR shows the highest sphericity variation of -7.9%. Hierarchical clustering groups RXR*α*/RAR*β* as most similar and GR as most distinct, reflecting their structural characteristics.

### Analysis of structure validation parameters

Comparison of Real-Space Correlation Coefficients (RSCC) values from experimental structures and pLDDT scores from AF2 structures revealed consistent patterns with overlapping scores in most NRs, particularly in their DBDs and LBDs (Fig. 4). Overall, AF2 predictions align well with experimental structures in well-structured regions like the DBDs and LBDs. However, notable deviations are observed in specific areas, such as the GR region near residue 616 (Fig. 4, GR red arrow), where all experimental structures show high RSCC scores, but the AF2 pLDDT score is low. Conversely, in the HNF4*α* 4IQR_B structure, for residue 145 (Fig. 4, HNF4*α* 4IQR_B purple arrow), AF2 shows pLDDT around 0.6, while RSCC drops to a negative value. For LXR*β*, regions with low RSCC values correspond to low-confidence pLDDT scores (pLDDT < 0.5). In NURR1 (7WNH_C, _D), RSCC values fluctuate significantly (0.61) around residue 300 (Fig. 4, NURR1 7WNH_C/D blue arc), but AF2 models maintain consistently high pLDDT scores (>0.9). PPAR*γ*, RAR*β*, and RXR*α* show consistent patterns of overlapping RSCC and pLDDT values, with localized drops in scores of varying magnitudes.

**Figure 4:**
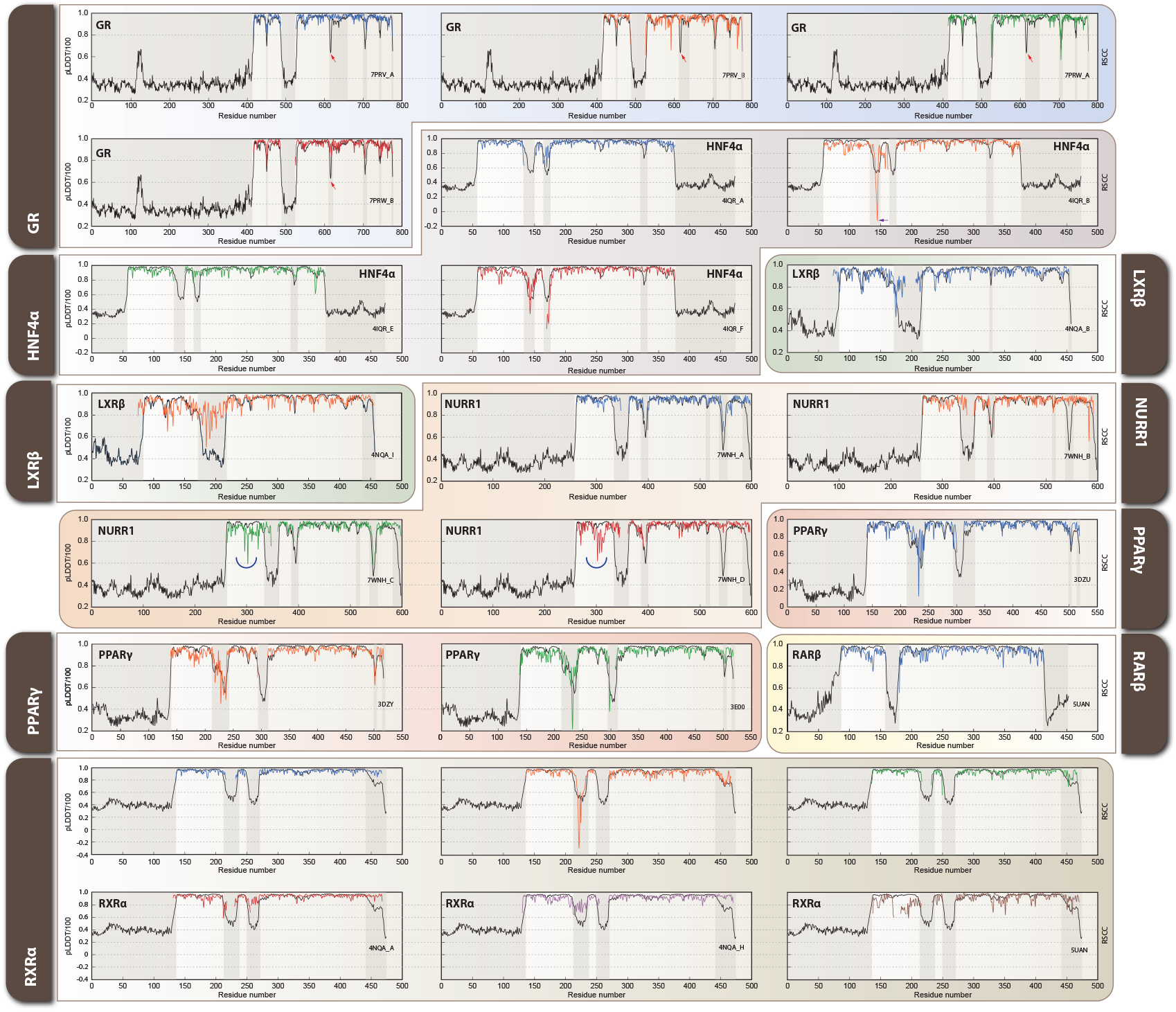
Real-Space Correlation Coefficients (RSCC) and pLDDT score comparison across analyzed NR. Black lines represent AF2 pLDDT scores, while colored lines show RSCC values from experimental structures: blue (first chain), orange (second chain), green (third chain), red (fourth chain), purple (fifth chain), and brown (sixth chain). Key regions of interest are highlighted: red arrow indicates GR residue 616, where experimental structures show high RSCC but AF2 predicts low confidence; purple arrow points to HNF4*α* 4IQR_B residue 150, where AF2 shows moderate confidence while experimental RSCC drops to negative values; blue arcs highlight NURR1 regions around residue 300 in chains C and D (7WNH), where experimental RSCC fluctuates significantly while AF2 maintains high confidence scores. These patterns reveal structure-prediction discrepancies that often correspond to functionally important flexible regions across the NR family.

Structural validation metrics RSCC, b-factors, and pLDDT were displayed in “putty” representation for DBD and LBD of illustrative NRs (Fig. 5). The tube diameter and color range are linked to structural confidence and flexibility. Narrow, blue tubes represent high-confidence structural metrics (high RSCC, low b-factors, and high pLDDT values), while large, red tubes indicate low-confidence regions. GR and RXR*α* (from RXR*α*-PPAR*γ* heterodimer, PDB: 3DZY) demonstrate high correlation between RSCC, b-factors, and pLDDT, as the “putty” representation is consistent across all shown metrics. Inversely, NURR1 and RXR*α* (from RXR*α*-LXR*α* heterodimer, PDB: 4NQA) exhibit lower conservation across structural validation metrics. Within GR DBD, many residues exhibit consistently high RSCC, low b-factors, and high pLDDT values, such as P418, K419, and L420, indicating strong structural agreement and stability in this domain. However, notable outliers include Q452 (Fig. 5 top panel) (RSCC = 0.74, b-factor = 101.99, pLDDT = 69.54), which deviates significantly from neighboring residues, suggesting a region of increased flexibility or a distinct structural feature. Similarly, N454 and H453 show elevated values, marking a transition to a more flexible region. In RXR*α*/PPAR*γ* complex the RXR*α* LBD, shows similar patterns that support structural stability and consistency, whereas NURR1 DBD and RXR*α* LBD in complex with LXR*β* exhibit cases where RSCC and pLDDT diverge, highlighting structurally ambiguous regions.

**Figure 5:**
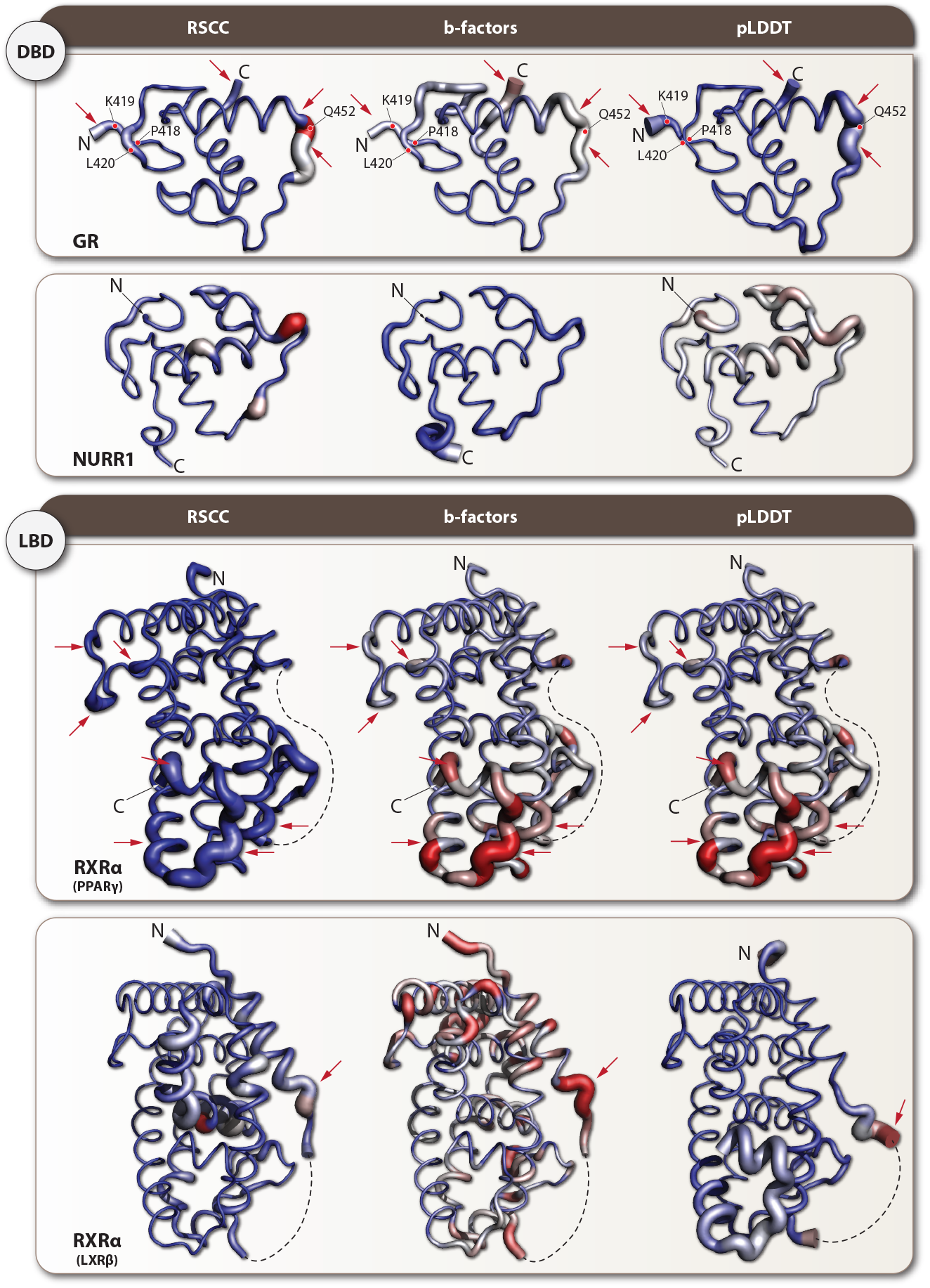
Structural validation metrics of illustrative NRs. “Putty” representation of GR DBD, NURR1 DBD, RXRRXR*α* as heterodimer of PPAR*γ* and in complex with LXR*β* highlights local structural confidence using RSCC, b-factors, and pLDDT data. Tube thickness and color intensity correspond to residue-level variability, with blue and narrow regions indicating high structural reliability (high RSCC, low b-factors, high pLDDT), whereas red and expanded regions mark flexible or uncertain areas. GR DBD shows consistently high confidence metrics for most residues (e.g., P418, K419, L420), with notable exceptions such as Q452 (RSCC = 0.74, b-factor = 101.99, pLDDT = 69.54), which represents a region of increased flexibility. High correlation across metrics is observed for GR and RXR*α* (PDB: 3DZY), where putty representations remain consistent across all validation metrics, indicating robust structural prediction. In contrast, NURR1 DBD and RXR*α* LBD in complex with LXR*β* (PDB: 4NQA) exhibit local discrepancies between experimental and computational validation metrics, particularly in regions where RSCC and pLDDT values diverge significantly, highlighting areas where AF2 predictions may not fully capture the conformational states observed in experimental structures.

### Backbone Torsional Analysis

We analyzed the backbone conformations between experimental crystal structures and AF2 predictions for seven NRs through Ramachandran plot distributions obtained from Molprobity validation server and quantitative metrics. The GR (7PRV_A) showed the highest structural agreement with *ϕ/ψ* correlations of 0.842/0.868 respectively, ranking first in overall correlation. The experimental structure contained 4 Ramachandran outliers while its AF2 counterpart showed only 1 outlier. In contrast, LXR*β* (4NQA_I) demonstrated the poorest agreement, with the lowest *ϕ/ψ* correlations (0.535/0.533) and highest *ψ* angle Wasserstein distance (13.99). The Ramachandran plots for these two representative cases, illustrating the best and worst agreement between experimental and AF2 structures, are shown in Fig. 6. HNF4*α* exhibited an interesting disparity between *ϕ* correlation (0.915, ranked 1st) and *ψ* correlation (0.751, ranked 16th). Quantitative comparison revealed systematic differences in stereochemical quality. The experimental PPAR*γ* structure (3E00_D) contained 18 Ramachandran outliers with 78.0% residues in favored regions, while its AF2 model showed only 1 outlier with 98.9% residues in favored regions. RAR*β* showed high stereochemical quality in both experimental (no outliers) and predicted structures. RXR*α* (3DZY_A) displayed high Wasserstein distances (10.62 for *ϕ*, 7.38 for *ψ*) despite moderate correlation values.

**Figure 6:**
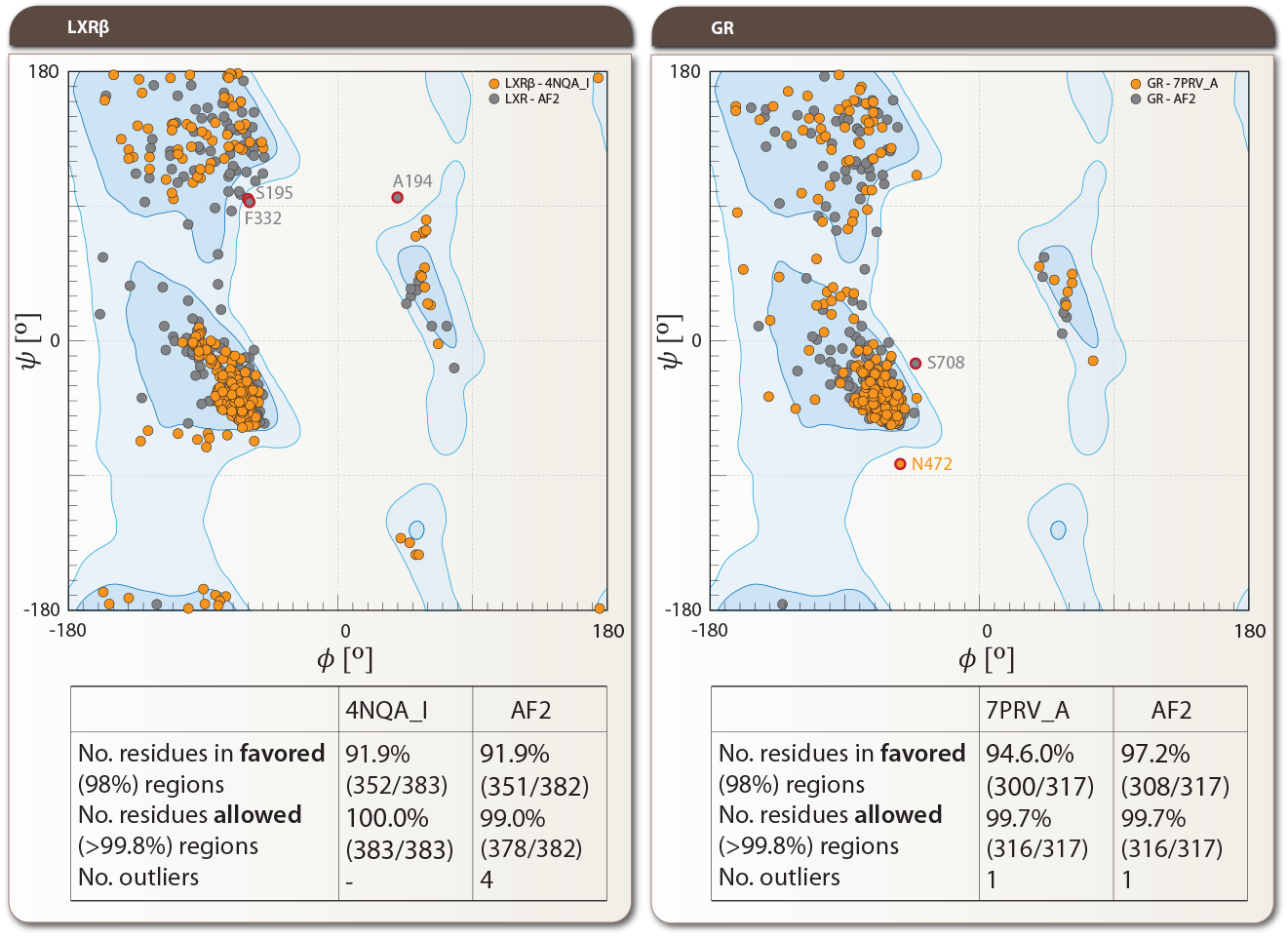
Ramachandran plot comparison between experimental structures and AF2 predictions for LXR*β* and GR. Left panels: LXR*β* 4NQA_I (experimental, orange circles) vs AF2 prediction (gray circles), showing the poorest agreement among analyzed NRs (*ϕ*/*ψ* correlations of 0.535/0.533 and Ψ angle Wasserstein distance of 13.99). Right panels: GR 7PRV_A vs AF2 prediction, demonstrating the highest structural agreement (*ϕ*/*ψ* correlations of 0.842/0.868) among the seven analyzed NRs. Specific outliers are labeled, including F332 and A194 for LXR*β*, and S708 and N472 for GR. Tables below each plot provide statistics highlighting the number of amino acids in favored (98%) and allowed (>99.8%) regions, as well as outliers. Note that the experimental GR structure contained 4 Ramachandran outliers while its AF2 counterpart showed only 1, reflecting AF2’s tendency toward higher stereochemical quality, potentially at the expense of functional conformational diversity. Data obtained from Molprobity validation server (http://molprobity.biochem.duke.edu).

## DISCUSSION

The observed RMSD patterns in NR predictions highlight both the strengths and challenges of AF2 in modeling multi-domain proteins. The generally low RMSD values (< 0.55 Å) for most structures align with previous benchmarking studies, which demonstrated AF2s high accuracy in predicting monomeric protein structures ^4^. This aligns with findings from cryoEM challenge datasets, where AF2 achieved consistent structural agreement between its models and cryoEM density maps. Analysis of 13 unique datasets showed that only one case required domain-level adjustments, with a clear correlation between pLDDT scores and model accuracy ^19^. However, the domain-specific variations observed, particularly in NURR1 and LXR*β*, suggest that certain NRs pose unique structural challenges. These discrepancies likely stem from factors such as domain flexibility, inter-domain interactions, or the absence of cofactors and ligands in the prediction pipeline, an issue that has been previously noted when modeling LBDs ^20^. The unexpected finding that 41.7% of cases exhibited higher RMSD values in DBDs compared to LBDs contrasts with the conventional notion that DBDs are generally more structurally conserved across the NR subfamily ^14,17^. One possible explanation is that AF2 predictions may not fully capture the rigid-body constraints imposed by DNA interactions in experimental structures. Additionally, the violation of the expected RMSD hierarchy (all-atom > backbone > C*α*) in 12.5% of cases reflects the inherent variability in AF2’s accuracy across different structural elements, particularly in flexible or solvent-exposed regions, as previously observed in protein structure prediction studies ^4,6,20,21^. Furthermore, while AF2 can predict multiple conformations, its accuracy varies depending on the protein’s flexibility and the presence of multiple domains. This aligns with our observation that AF2’s predictions show variability in flexible regions, such as LBDs, leading to deviations in the expected RMSD hierarchy. Therefore, the challenges in predicting flexible or solvent-exposed regions may contribute to the observed discrepancies in our study ^22^. Our observation of domain-specific variations, particularly pronounced in NURR1 (CV = 0.7% for DBD vs. 18.0% for LBD), supports previous findings that AF2’s performance can be influenced by missing cofactors and allosteric effects ^6^. Structural variability in LBDs is expected given their functional plasticity, which is required for ligand-binding and conformational switching. The results suggest that while AF2 provides highly accurate backbone predictions, domain-specific challenges remain, particularly for proteins with dynamic regulatory elements.

The systematic differences observed in SSE predictions, particularly the tendency toward higher *α*-helical content in AF2 models, align with previous observations by Jumper *et al*. ^4^ and Varadi *et al*. ^6^. The strong negative correlation between *α*-helical and loop content (r = -0.906 for LBD) suggests a compensatory mechanism in structural prediction, possibly reflecting the algorithm’s training data composition. The complete absence of *β*-sheets in some experimental RXR*α* structures, while present in AF2 predictions, corresponds with previous findings regarding the challenges in predicting less prevalent SSEs ^20^. This discrepancy likely stems from the statistical nature of the deep learning approach, where AF2 may favor predicting more commonly observed structural elements in the training data to minimize overall prediction error. Less frequently occurring structural features, such as the absence of *β*-sheets in certain conformational states, may be suppressed in the model’s predictions as they represent statistical outliers in the training dataset. In addition, this disparity might arise from the inherent flexibility of these regions in solution, as demonstrated by Nuclear Magnetic Resonance (NMR) and hydrogen/deuterium exchange (HDX) studies ^23^, which may not be fully captured in either experimental 3D structures or AF2 predictions. The observed variations in domain-specific prediction accuracy suggest that local sequence context and domain interactions play crucial roles in determining secondary structure assignment. This is further supported by studies comparing crystal and cryo-EM structures of P-loop channels with AF2 models, which showed strong overall similarity but notable differences in *π*-bulge patterns ^24^.

The geometric analysis of NR domain organization reveals complex patterns of structural variability and domain positioning that reflect both intrinsic flexibility and partner-specific adaptations. Our analyses focused on the DBD and LBD domains, as the intervening hinge region is typically absent from crystal structures due to its inherently flexible and disordered nature. The observed variations in domain distances, angles, and dihedrals demonstrate that NRs can adopt multiple stable conformations, consistent with their allosteric regulation mechanisms ^25^. The most pronounced irregularities were observed in the inter-domain distances of HNF4*α* and NURR1, which showed the largest interquartile range (IQR) values among all analyzed receptors. This bimodal distribution of conformations aligns with recent findings by Chandra *et al*. ^18^, who demonstrated that NRs can exist in multiple distinct conformational states depending on their functional context. The substantial variation in HNF4*α* domain separation (IQR = 27.72 Å) particularly reflects its documented ability to form asymmetric dimers, where one monomer adopts an extended conformation while its partner maintains a more compact arrangement ^26^. Domain angles showed receptor-specific patterns of variability, with RXR*α* exhibiting particularly interesting partner-dependent conformations. This observation supports RXRs’ adaptabilities, as RXR*α*’s structural plasticity allows it to adjust to different heterodimeric partners ^23,27^. Notably, our analysis reveals a distinct pattern in conformational flexibility between homodimerizing and heterodimerizing NRs. Receptors capable of homodimerization demonstrate greater conformational diversity, particularly evident in DNA-bound structures where one monomer adopts a compact conformation while its partner maintains an extended state. This asymmetric organization, observed in both GR and HNF4*α* structures, likely reflects their need to accommodate symmetric self-association and represents a fundamental mechanism for DNA binding. In contrast, NRs that exclusively heterodimerize with RXR*α* show more constrained conformational ranges, suggesting evolutionary optimization for specific partner interactions. RXR*α* itself exhibits an intermediate level of flexibility that allows it to serve as a promiscuous dimerization partner. Interestingly, AF2 predictions capture only one conformational state, typically matching the more extended conformation, highlighting the importance of experimental structures in revealing the full spectrum of biologically relevant conformational states. The systematic differences in RXR*α* angles when partnered with PPAR*γ* versus LXR*β* (∼50° difference) align with recent cryo-EM studies showing partner-specific conformational selection in NR heterodimers ^15,17^. The dihedral angle distributions revealed unexpected patterns, particularly in NURR1 and GR structures, where distinct conformational clusters were observed. Recent molecular dynamics studies that suggest that such conformational diversity might be essential for allowing these receptors to interact with different regulatory elements and coregulators ^28,29^. The bimodal distribution of GR dihedral angles (IQR = 81.81°) particularly reflects its documented ability to adopt distinct conformations in response to different ligands ^30,31^. AF2 predictions generally captured the predominant conformational states observed in experimental structures, but showed interesting deviations in cases of extreme structural plasticity. This aligns with the assessments of AF2’s performance on flexible multi-domain proteins ^20^, suggesting that while the algorithm effectively predicts stable conformations, it may not capture the full range of physiologically relevant structural states. The boxplot analysis (Fig.2A-C) highlights key patterns in NR structural variability. HNF4*α* and NURR1 exhibit greater conformational diversity, likely reflecting their functional plasticity. RXR*α* shows partner-dependent clustering, supporting the idea that its conformation is influenced by heterodimeric interactions ^32^. Domain-specific differences reveal that DBD-LBD distances are more variable than angles, suggesting inter-domain separation plays a key role in NR allostery. A more detailed structural comparisons revealed receptor-specific patterns: i) AF2’s HNF4*α* structure shows greater similarity to specific chains (4IQR_B, _F) that facilitate high-affinity DNA-binding ^14^, ii) the LXR*β* model closely matches specific conformational states observed in crystal structures (4NQA_I), particularly in terms of domain geometry parameters and iii) RXR*α*’s predicted structure shows variable similarity to different experimental structures depending on the geometric descriptor analyzed, consistent with its known adaptability to different dimerization partners ^18^.

The high accuracy in sphericity predictions, with r=0.891, indicates that AF2 captures the overall pocket geometry well, consistent with its demonstrated ability to predict global protein structure ^4^, whereas there was a negative correlation between effective pocket radius and sphericity with r=0.482. Notably, RXR*α* shows exceptional consistency between experimental and predicted structures, potentially due to its well-conserved pocket architecture ^25^. The larger LBP volume range in LXR*β* (−15.8%) aligns with studies highlighting its structural flexibility ^33^, suggesting that static AF2 predictions may not fully capture its dynamic LBP. Moreover, the systematic underestimation of pocket volumes by AF2 suggests that the algorithm may be biased toward capturing more compact, energetically favorable states rather than the full range of physiologically relevant conformations that NRs adopt during transcriptional regulation. An additional study comparing AF2-predicted and experimental structures of G-protein-coupled receptors (GPCRs) provides further evidence of these limitations ^34^. While AF2 can predict protein structures with a C*α* RMSD accuracy of 1 Å, it struggles with high-accuracy predictions of GPCR pocket side chains. It also poorly predicts extracellular and transmembrane domains, as well as the transducer-binding interface of GPCRs. These findings suggest that AF2 may be less reliable for modeling highly flexible or functionally dynamic regions, reinforcing the importance of experimental structure determination for capturing biologically relevant conformations ^34^. All these findings expand our understanding of AF2’s capabilities and limitations in predicting NR binding pockets, which is particularly important for structure-based drug design applications ^35^.

The comparison of structure validation parameters between AF2 predictions and experimental structures showed distinct patterns in NR conformational stability. Differences between RSCC and pLDDT scores are more pronounced in loops or unstructured regions, such as the activation function-1 domain at the N-termini and certain C-terminal regions, where experimental structures are often unavailable. The activation function-1 domain is particularly notable for its uniformly low RSCC and pLDDT values, reflecting the lack of experimental structural data and its inherent flexibility. These findings highlight both the strengths of AF2 predictions in core domains and the limitations in capturing certain unstructured or flexible regions. This observation is supported by a comprehensive analysis of 904 human proteins comparing AF2 and NMR structures, which found that while AF2 models are generally more accurate than NMR structures, NMR structures prove more reliable for dynamic structures ^36^.

For GR residue S616, located in helix H5 of the LBD, experimental structures consistently show high RSCC values while AF2 predicts low confidence. This region forms part of the LBD-DBD interface that is critical for receptor function - specifically, the N-terminal end of H5 forms substantial interactions with DBD and participates in key domain interfaces when bound to ligands like velsecorat or fluticasone furoate ^13^. The high RSCC values in crystal structures suggest this region becomes well-ordered in the presence of binding partners, while AF2’s low confidence may reflect inherent flexibility in the unbound state. For HNF4*α* residue E145 in the hinge region close to helix H1, AF2 predicts moderate confidence (pLDDT > 0.6) while experimental structures show very low or negative RSCC values specifically in chains B and F of the 4IQR structure. This region is part of a critical DBD-hinge-LBD construct (residues 141-368) that facilitates high-affinity DNA binding, as directly demonstrated by the 75-fold weaker binding of isolated DBD-hinge portions (*K*_*d*_ ∼6000 nM) compared to constructs including the LBD (*K*_*d*_ = 80 nM) ^14^. The negative RSCC values in specific chains suggest conformational heterogeneity at these interfaces rather than inherent disorder, consistent with their role in dynamic domain organization required for cooperative DNA binding. These discrepancies between computational predictions and experimental structures highlight regions involved in conformational changes and allosteric regulation, providing valuable insights for understanding NR function and potential drug targeting strategies. For all NRs, the low pLDDT scores in the AF2 correlate with the missing residues or low RSCC values in experimental 3D structures. As expected, pLDDT scores also inversely correlate with b-factor values. This correlation between pLDDT scores and RSCC values has been independently validated in a large-scale comparison of 150,000 human protein structures, though that study also emphasized that high-resolution crystallographic structures remain preferable when available ^37^. Notably, several NRs (PPAR*γ*, RXR*α*, HNF4*α*) exhibit missing residues in their LBD regions, and these areas consistently show lower pLDDT scores (50-70) in AF2 predictions, even when other experimental structures contain these regions. Conversely, NRs with complete experimental structures (RAR*β*, LXR*β*) show consistently high pLDDT scores (>90) throughout their LBDs. Interestingly, for NRs lacking crystal structures (ROR*β*, GCNF, NOR1, TR2, EAR2), AF2 generally predicts high confidence scores (>70-90) across their LBDs, with only occasional lower-confidence regions.

Our backbone torsional analysis reveals a consistent pattern where AF2 models demonstrate higher stereochemical quality compared to experimental structures. This observation aligns with that AF2’s energy function strongly favors ideal backbone geometry and AF2’s architecture implements strong geometric constraints during structure prediction, explaining the high proportion of residues in favored Ramachandran regions ^4^. The “imperfections” observed in crystal structures likely reflect genuine conformational flexibility or structural adaptations supported by finding that crystal structure Ramachandran outliers often occur in functionally relevant regions and may represent legitimate conformational states rather than experimental errors ^38^. The varying degrees of agreement between experimental and predicted structures across different NRs suggest that AF2’s accuracy depends on multiple factors. Similar observations were reported by Akdel *et al*. ^20^ in their large-scale analysis of AF2 predictions, particularly noting that regions of proteins involved in interactions or conformational changes may show larger deviations from idealized geometry. The case of HNF4*α*, with its disparate *ϕ*/*ψ* correlations, illustrates how AF2 may capture overall fold topology while missing specific local conformational details. This aligns with findings showing that while AF2 excels at predicting global structure, it may miss subtle but functionally important local conformational features ^39,40^. The high-quality predictions for RAR*β* demonstrate that AF2 can achieve exceptional accuracy when the native structure already exhibits ideal geometry. However, the systematic differences observed in other cases raise important considerations for structure-based drug design and protein engineering applications, where natural conformational flexibility may be functionally relevant as shown for understanding enzyme mechanisms and designing effective inhibitors ^41^. While AF2 has shown remarkable potential in predicting individual protein structures, it’s important to note that proteins rarely function in isolation. Recent developments in AF2 and in particular in AF3 have begun to address the challenges of modeling macromolecular complexes, though experimental methods remain essential for validation and for capturing the full range of biological conformations ^19,42^.

To provide a comprehensive overview of our comparative analysis between experimental and AF2-predicted NR structures, we summarize the eight key metrics evaluated in this study in Table 1. This summary highlights both the strengths and limitations of AF2 in capturing the structural features critical for NR function.

## KEY LIMITATION OF AF2 IN NR PREDICTION AND FUTURE DIRECTIONS

The comprehensive analysis of AF2 predictions for NRs shows several key limitations in current structure prediction approaches. A fundamental challenge lies in AF2’s tendency to model static, well-ordered conformations that mirror crystallographic states, potentially missing important physiologically relevant conformational heterogeneity. Current limitations can be categorized into three main areas:

1. The first major limitation concerns dynamic regions and conformational flexibility. Our analysis showed significantly higher structural variability in LBDs (CV = 29.3%) compared to DBDs (CV = 17.7%), with systematic underestimation of LBP volumes. This is particularly evident in cases like LXR*β*, where we observed a -15.8% deviation in pocket volume predictions. This limitation is particularly significant for structure-based drug design approaches targeting NRs, where accurate pocket volume and flexibility prediction is essential for identifying potential ligands.
2. The second key limitation involves multi-domain organization and regulation. AF2 shows notable difficulties in capturing multiple biologically relevant conformational states, particularly the compact-extended asymmetry in homodimers and predicting partner-specific conformational adaptations in heterodimeric complexes.This limitation likely originates from the algorithm’s optimization for single-state prediction rather than conformational ensembles, and its reduced capacity to model the cooperative domain movements and long-range allosteric effects that are crucial for NR function. Our analysis revealed that AF2 consistently predicts only one conformational state for asymmetric homodimers like GR and HNF4*α*, typically corresponding to the extended conformation, while missing the biologically significant compact states that enable differential DNA recognition and cofactor recruitment.
3. The third limitation stems from training biases toward crystallizable protein states and static structural data, affecting the overall prediction quality particularly in flexible regions. This bias is compounded by the underrepresentation of multiple conformational states for the same protein in structural databases, where proteins are typically captured in single, stable conformations favored by crystallization. Additionally, the algorithm may have inherent difficulties in capturing the long-range interactions that govern domain-domain positioning across different functional states, particularly when these relationships involve subtle allosteric mechanisms or partner-induced conformational changes. This limitation particularly affects NRs, where conformational diversity is essential for their function as molecular switches in transcriptional regulation.

To address these challenges, the integration of molecular dynamics simulations and energy minimization protocols could significantly improve conformational landscape exploration, though this would increase computational costs. Expanding training datasets to include diverse experimental data from cryo-EM and NMR studies could enhance the prediction of dynamic states and transient conformations.

The future of NR structure prediction likely lies in hybrid approaches that integrate experimental and computational methods. While more sophisticated sampling methods and broader training datasets could enhance prediction accuracy, they must balance computational efficiency with biological relevance. Future methods should focus on capturing both structural accuracy and complex regulatory mechanisms, including domain flexibility, allosteric communication, and partner-specific adaptations. Given AF3’s expanded capabilities in predicting protein complexes, systematic evaluation following our analysis framework is needed, particularly focusing on DNA binding, ligand interactions, and cofactor peptide associations in NRs.

## CONCLUSION

This first comprehensive comparative analysis of AF2-predicted and experimental NR structures reveals both the strengths and limitations of computational structure prediction. While AF2 excels at predicting stable conformations with high stereochemical quality, it shows limitations in capturing the full spectrum of biologically relevant conformational states, particularly in flexible regions and LBPs. The study demonstrates that AF2 predictions should be used as complementary tools alongside experimental structures, especially for understanding protein dynamics and drug-target interactions. Critical differences between predicted and experimental structures often highlight functionally important regions involved in allosteric regulation and protein-protein interactions, providing valuable insights for structure-based drug design targeting NRs.

## MATERIALS AND METHODS

All geometrical measurements and analysis have been done in PyMOL v2.4.2 ^43^ using internal commands or python scripts.

### RMSD analysis

The RMSD is a global measure of the similarity between protein structures, calculated as the square root of the mean squared distances between aligned atoms of superimposed structures. The structure-based dynamic programming alignment of experimental and predicted structures was determined using three methods: i) all-atoms, ii) only C*α* carbons and iii) backbone (N, C*α*, C and O atoms). The RMSD in Å was calculated individually for the DBDs and LBDs using the `super` command in PyMOL (Version 2.5, Schrödinger, LLC) according to the equation:

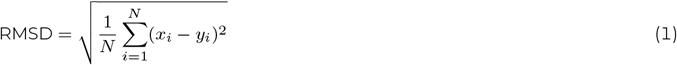

where *N* is the number of aligned atoms, and (*x*_*i*_ − *y*_*i*_) represents the distance between the *i*th pair of equivalent atoms after optimal superposition. Statistical analyses were performed to quantify structural variations and domain-specific differences. The coefficient of variation (CV) was calculated to assess relative variability:

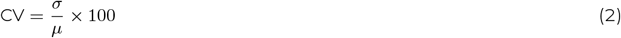

where *σ* is the standard deviation and *μ* is the mean RMSD value for each measurement type. Domain-specific differences were evaluated using paired statistical analysis, with effect size quantified by Cohen’s *d*:

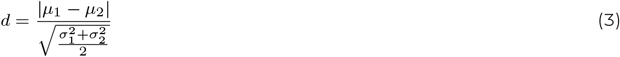

where *μ*_1_ and *μ*_2_ are the means, and *σ*_1_ and *σ*_2_ are the standard deviations of the DBD and LBD RMSD values, respectively. The significance of domain differences was assessed using a paired t-test with a significance level of *α* = 0.05. To assess structural prediction consistency across different measurement types (all-atom, backbone, C*α*), hierarchical patterns were analyzed, and deviations from the expected pattern (all-atom > backbone > C*α*) were quantified. Domain measurements were performed separately to account for the potential positional bias arising from differential domain arrangements in 3D space.

### SSE content analysis

The *α*-helical, *β*-sheet, and loop content expressed in % was calculated separately for DBD and LBD using python scripts in PyMOL environment. Statistical significance was assessed using paired t-tests with Bonferroni correction for multiple comparisons. Structure element correlations were calculated using Pearson’s correlation coefficient:

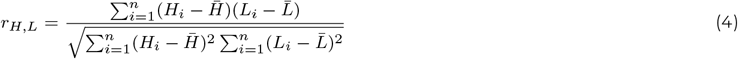

where *H*_*i*_ and *L*_*i*_ represent *α*-helical and loop content percentages, respectively.

### Geometric analysis of domain architecture

#### Distance *ϵ*_*DBD,LBD*_ between DBD and LBD

The physical distance in Å between the two domains (DBD and LBD) was calculated by internal command `get_distance` between the two COMs located in two domains using the `centerofmass.py` script. Statistical variation in domain distances was quantified using the interquartile range (IQR):

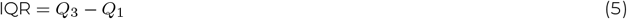

where *Q*_1_ and *Q*_3_ represent the first and third quartiles of the experimental structure measurements, respectively.

#### Angle *θ*_*DBD,LBD*_ between DBD and LBD

*θ*_*DBD,LBD*_ was calculated using three points 1) DBD_*COM*_, 2) LBD_*COM*_, and the 3) COM defined by using the last C*α* atom of DBD (DBD_*CαL*_) and the first C*α* atom of LBD (LBD_*CαF*_). This approach was chosen to standardize the definition of the hinge COM across structures with varying numbers of missing atoms in the hinge region. Angles are expressed in degree (°) and were calculated using the PyMOL command `get_angle`.

Angular variation was assessed using both IQR and circular statistics:

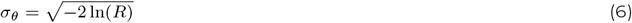

where *R* is the mean resultant length of the angular measurements.

#### Dihedral angles *DH*_*DBD,LBD*_ between DBD and LBD

*DH*_*DBD,LBD*_ were calculated using four points: 1) DBD_*COM*_, 2) DBD_*CαL*_, 3) LBD_*CαF*_ and 4) LBD_*COM*_. Angles are expressed in degrees and were calculated using the PyMOL command `get_dihedral`.

Dihedral angle distributions were analyzed using circular statistics and conformational clustering:

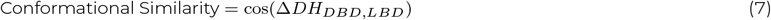

where Δ*DH*_*DBD,LBD*_ represents the difference in dihedral angles between structures.

Comparative analysis between experimental structures and AF2 predictions was performed using multiple metrics. The spread of experimental measurements was quantified using IQR to assess structural variability. Partner-specific effects were evaluated using one-way ANOVA with post-hoc Tukey’s test where appropriate. For circular measurements (angles and dihedrals), appropriate circular statistics were applied. Statistical significance was set at p < 0.05.

### Analysis of LBP geometry

The LBPs were identified by hollow 1.3 ^44^. The Connoly surface volume and are were calculated with 1.4 Å probe radius using msms 2.6.1^45^ and the PyMOL command `get_area`, respectively. The LBPs were subsequently visualised by PyMOL at cutoff value of 5 Å from the particular ligand located in the structure. For the *apo* AF2 predicted structure the cutoff was calculated from the position of all merged ligands in the *holo* structures. The LBP sphericity Ψ, is defined as:

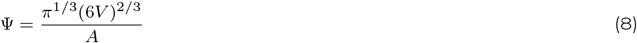

where is *V* is the volume and *A* is the surface area. The effective radius of the LBP is defined as the radius of a sphere with same surface area to volume ratio as the volume of interest, determined by:

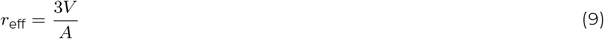

Statistical analyses for the LBP geometry were performed using custom Python scripts. Kruskal-Wallis H-tests were conducted to assess significant differences between NRs for both effective radius and sphericity measurements. Pearson correlation coefficients were calculated to evaluate relationships between effective radius, sphericity, and experimental versus AF2 predicted values. For experimental structures with multiple measurements, means and standard deviations were calculated. Hierarchical clustering was performed using Euclidean distances between radius and sphericity measurements to group receptors with similar characteristics. Linear regression analysis was used to assess the systematic relationships between experimental and AF2-predicted values, with regression slopes indicating prediction biases. Percentage deviations between experimental and predicted values were calculated as (*v*^pred^ − *v*_exp_)*/v*_exp_ × 100%, where *v* represents either radius or sphericity measurements.

### Analysis of structure validation parameters

#### Comparative analysis of b-factors, RSCC and pLDDT parameters

Local structural quality metrics were visualized through spatial mapping using PyMOL’s b-factor putty representation. The visualization parameters were configured to display variations in putty diameter and chromatic gradient corresponding to the respective quality metrics. For crystallographic structures, atomic b-factors were utilized to assess conformational mobility, while RSCC provided quantitative measures of model-to-density agreement. pLDDT scores derived from AF2 models served as confidence metrics for the computational predictions. To enable direct comparison, pLDDT values were scaled by 1/100 to match the same numerical range as RSCC. RSCC of PDB structures and pLDDT scores of AF2 structures were then compared and plotted using AF2 NR sequence numbering. Comparative analysis of these spatially mapped parameters facilitated the identification of regions exhibiting structural correspondence or deviation between experimental and computationally predicted models, with particular emphasis on the DBD and LBD regions.

#### Torsional Analysis

The backbone torsion angles (*ϕ* and *ψ*) of the DBDs and LBDs were analyzed using a custom Python script implemented through the PyMOL molecular visualization system. Based on *ϕ* and *ψ* torsion angles, the similarity between experimental structures and AF2 predictions was assessed using Python script. Four metrics (RMSD, Wasserstein Distance, Mean Absolute Error, Pearson Correlation Coefficient) were used to evaluate the resemblance for each NR pair. Conformational analysis of the protein backbone geometry was performed using Ramachandran plot calculations in MolProbity ^38^ hosted at http://molprobity.biochem.duke.edu website. Prior to analysis, structural alignment between AF2-predicted models and experimentally determined structures was performed to ensure proper comparative assessment. The Ramachandran plots were generated by submitting each NR structure to the MolProbity server, enabling quantitative evaluation of backbone conformations through assessment of sterically allowed and disallowed regions in the *ϕ* - *ψ* torsional space.

## ACKNOWLEDGMENTS

This work was supported by Nazarbayev University Collaborative Research Proposal # 091019CRP2108 to F.M.

## AUTHOR CONTRIBUTIONS

**Akerke Mazhibiyeva**: Methodology, Investigation, Formal analysis, Visualization & Writing the original draft.

**Tri P. Pham**: Formal analysis, Writing & Editing.

**Karina Pats**: Formal analysis, Writing & Editing.

**Martin Lukac**: Formal analysis, Writing & Editing.

**Ferdinand Molnár**: Conceptualization, Funding acquisition, Supervision, Visualization, Writing & Editing.

